# *k*-nonical space: sketching with reverse complements

**DOI:** 10.1101/2024.01.25.577301

**Authors:** Guillaume Marçais, C.S. Elder, Carl Kingsford

## Abstract

Sequences equivalent to their reverse complements (i.e., double-stranded DNA) have no analogue in text analysis and non-biological string algorithms. Despite this striking difference, algorithms designed for computational biology (e.g., sketching algorithms) are designed and tested in the same way as classical string algorithms. Then, as a post-processing step, these algorithms are adapted to work with genomic sequences by folding a *k*-mer and its reverse complement into a single sequence: the canonical representation (*k*-nonical space). The effect of using the canonical representation with sketching methods is understudied and not understood. As a first step, we use context-free sketching methods to illustrate the potentially detrimental effects of using canonical *k*-mers with string algorithms not designed to accommodate for them. In particular, we show that large stretches of the genome (“sketching deserts”) are undersampled or entirely skipped by context-free sketching methods, effectively making these genomic regions invisible to subsequent algorithms using these sketches. We provide empirical data showing these effects and develop a theoretical framework explaining the appearance of sketching deserts. Finally, we propose two schemes to accommodate for these effects: (1) a new procedure that adapts existing sketching methods to *k*-nonical space and (2) an optimization procedure to directly design new sketching methods for *k*-nonical space.

The code used in this analysis is freely available at https://github.com/Kingsford-Group/mdsscope.

## 1 Introduction

Genomics sequence analysis shares many similarities with text analysis since they are both concerned with efficiently storing and searching long strings. Consequently, many algorithms are common between the two fields. The double-stranded nature of DNA brings a unique twist to genomics sequence analysis: a sequence and its reverse complement, in many cases, should be considered identical. For example, a sequencing read can represent either strand of a chromosome, and a read aligner attempts to find the best alignments of the read and its reverse complement against a genome. This concept has no equivalent in text analysis.

Sketching methods (e.g., minimizers [19, 20], syncmers [4]) create a small representation of a sequence (a “sketch”) by selecting a subset of *k*-mers (substrings of fixed length *k*) from the original sequence. Because of their small size, sketches allow for efficient sequence similarity estimation that is much faster than comparing the original sequences. Thus, sketching is a common strategy to make bioinformatics algorithms more efficient (see [14] and [26] for reviews). To handle reverse complements, sketches usually work with “canonical *k*-mers,” i.e., the smallest (lexicographically) of *m* and the reverse-complement 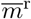. By using this canonical representation, denoted by *m*^c^, a *k*-mer and its reverse complement are treated equally.

More precisely, for our purposes, a sketching method is a function *φ* that takes as input one or more *k*-mers (the context) and outputs a (possibly empty) set of the indices of the *k*-mers to select. A sketch of a sequence *S* is constructed by collecting all the *k*-mers selected by *φ* over all the contexts of *S*. Sketching methods are usually designed and evaluated as string algorithms working on standard strings. Then, to handle reverse complements, implementations of sketching methods do not use *φ* directly but rather work in “*k*-nonical space”: the selection function is composed with the canonical function, i.e., instead of querying *φ*(*m*) one queries *φ*(*m*^c^). See for examples the minimap [9, 10] aligner using minimizers and the modified version using parameterized syncmers [3].

There exist many sketching functions, and their performance has been studied in diverse settings; however, the effect of using canonical *k*-mers has received little attention. This is surprising given the importance of reverse complements to computational biology. Moving to *k*-nonical space has been primarily viewed as an implementation detail that is not likely to have an impact on downstream applications. Unfortunately, as we show, moving to canonical space can have a significant impact.

An important property of sketching methods is the *window guarantee*, i.e., the assurance that the distance between two selected *k*-mers in an input sequence *S* is not too large. There are nuances to what this guarantee entails (e.g., if it is probabalistic, if some subsequences are excluded, etc.) that we discuss in Section 2.2. In essence, it means that *k*-mers are selected at approximately regular intervals from an input sequence and that no large regions of the genome are ignored. Subsequent algorithms that use sketching typically require the window guarantee to prove their correctness.

As we show in Section 3, using a sketching method in canonical space may not preserve the original method ‘s window guarantee. This creates “sketching deserts,” i.e., regions of the genome where few or no *k*-mers will be selected. Consequently, these regions are ignored or under-represented in subsequent analysis (e.g., missing alignments), which creates unexpected statistical biases.

We first look at the structure of the de Bruijn graph and show how the window guarantee corresponds to strongly connected components (SCCs) of this graph. This develops the theory of how moving to canonical space can negate the window guarantee of a sketching method. Then we show the effect of canonical space in creating sketching deserts for common context-free sketching methods (methods taking as input exactly one *k*-mer) in a realistic scenario.

We propose two alternative strategies. First, we introduce the canonicalization optimization problem that transforms an existing sketching method into one that is “symmetric” (i.e., handles a *k*-mer and its reverse complement identically) while respecting the same window guarantee as the original method and not introducing any additional sketching deserts. Second, we give an ILP construction of context-free sketching methods that are symmetric by design and satisfy a prescribed window guarantee.

As seen in Section 3.3, there is an asymmetry between the sketching methods with a context used in practice (e.g., minimizers) and the context-free methods (e.g., syncmers). Because minimizers always select a *k*-mer in every context, it has the same window guarantee before and after canonicalization and is therefore immune to the detrimental effects of canonicalization described here. By contrast, every context-free method is susceptible to not having any window guarantee in *k*-nonical space. This asymmetry may partially explains why this effect was not recognized for context-free methods. In general, the effect of *k*-nonical space on the window guarantee is hard to predict, so sketching methods must be designed and tested to be robust to canonical representations before being used with genomics data.

## 2 Preliminaries

### 2.1 Notations

All sequences are strings over the alphabet ∑ of size *σ* = |∑|, and ∑ = {0, …, *σ −*1}. For the sake of simplicity, we assume that *σ* is even (*σ* = 2 or *σ* = 4 in our examples). Moreover, every letter of the alphabet has a complement: *a, b* ∈ ∑ are complements of each other when *a* + *b* = *σ−* 1. This is denoted as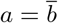 or (equivalently)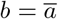. The genomic case corresponds to the mapping `A` = 0, `C` = 1, `G` = 2, `T` = 3. A sequence is an element of σ^*\**^ and *S*[*i* : *c*] is the substring of *S* starting at offset *i* and of length *c*.

Given a *k*-mer *m* = *m*_1_ … *m*_*k*_, its reverse complement is the *k*-mer 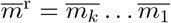. The set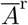contains the reverse complemented *k*-mers of set *A*. The *canonical representation* of *m* is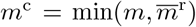, i.e., the lexicographically smallest of *m* and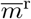. The sets 𝒞 and 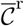denote canonical *k*-mers and their reverse complements, respectively. These sets cover the *k*-mers (i.e., 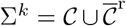) but may not be disjoint. Specifically, when *k* is even, the intersection 𝒞∩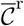 is the set of self-reverse complement *k*-mers—e.g., *k*-mers such as 0011 the intersection in the binary alphabet.

In general, a sketching method is a function *φ* : ∑^*k*^× · · · × ∑^*k*^→ 𝒫 ([1, *c*]). That is, the sketching function takes *c k*-mers as input (the context) and returns a possibly empty list of *k*-mers to select from this context. The context of length *c* of *S* at position *i* is the list of the *c* consecutive *k*-mers starting at position *i*: C(*S, c, i*) = [*S*[*i* : *k*], …, *S*[*i* + *c −* 1 : *k*]]. The sketch associated with φ for an input sequence *S* is the set of the offsets of all selected *k*-mers from every context of *S*: ℳ_*φ*_(*S, c*) = ∪_*i*_{*i* + *o* − 1 | *o* ∈ *φ*(C(*S, c, i*))}.

This general sketching definition can model methods with context (where *c >* 1, e.g., minimizers [19, 20] and minmers [8]), and context-free methods (where *c* = 1, e.g., syncmers [4, 3], fractional minimizers [21]). We focus on context-free methods where the input to the selection function is a single *k*-mer, and the output is a binary decision: do or do not select this *k*-mer. A context-free sketching selection function is defined by the set of all *k*-mers that it selects among all possible *σ*^*k*^ *k*-mers: P_*φ*_ = {*m* ∈ ∑^*k*^ | *φ*(*m*) ≠ ∅}. Equivalently, the function *φ* is the indicator function of the set P_*φ*_. Thus, computing the sketch reduces to computing the intersection between the *k*-mers in the sequence and the set P_*φ*_: ℳ_*φ*_(*S*, 1) = {*i* ∈ [0, |*S*|*−k* +1] | *S*[*i* : *k*] ∈ P_*φ*_}. Going forward, we do not differentiate between the selection function *φ* and the set P_*φ*_, and we freely use *φ* with set notations.

### 2.2 Cycles of the de Bruijn graph and selection deserts

A scheme *φ* has a *strong window guarantee* of *w* if the maximum distance between any two consecutively selected *k*-mers in any sequence is at most *w*. A scheme has a *relaxed window guarantee* if there exists only a small number of well-characterized repetitive sequences not intersecting *φ*. For example, the low entropy sequences {`AA` …, `CC` …, `GG` …, `TT` … } may be used as a set of non-intersecting sequences. When neither holds, a scheme has no window guarantee.

These notions have graph theoretical equivalents using the de Bruijn graph. The de Bruijn graph B_*k*_ of order *k* is the directed graph with *σ*^*k*^ nodes (one for each distinct *k*-mer) and an edge *u→ v* when the (*k−* 1)-suffix of *u* is equal to the (*k−* 1)-prefix of *v*. There is a one-to-one correspondence between the sequences of ∑* and the walks in B_*k*_. Moreover, a cycle in the de Bruijn graph corresponds to an infinitely long repetitive sequence.

An equivalent definition of the strong window guarantee is that the graph B_*k*_ *∖φ* (the de Bruijn graph with the *k*-mers of P_*φ*_ removed) is a directed acyclic graph (DAG) and the longest path in this DAG is of length *≤ w*. In that case, *φ* is called a decycling set of the de Bruijn graph. When B_*k*_ *∖φ* has strongly connected components (i.e., it contains cycles), *φ* does not have a window guarantee for any *w*. Similarly, *φ* has a relaxed window guarantee if the strongly connected components (SCCs) contain only cycles of sequences that we are willing to ignore (e.g., the low entropy sequences shown previously). For a function *φ*, SCC(*φ*) denotes the union of all the strongly connected components of B_*k*_\*φ*.

Given a *k*-mer *m* = *m*_0_*m*_1_ … *m*_*k−*1_, its rotation *k*-mer is R(*m*) = *m*_1_ … *m*_*k−*1_*m*_0_, and there is an edge *m→* R(*m*) in the de Bruijn graph. The cycle of all the rotations of *m* (*m*, R(*m*), R(R(*m*)), …) is called, for historical reasons, a Pure Cycling Register (PCR). The PCRs partition the nodes of the de Bruijn graph and play a special role in describing decycling sets. In particular, a decycling set must contain one node from each PCR and the minimum size decycling sets (MDSs) contain exactly one node from each PCR. Note that the converse is not true: not all sets containing exactly one node from each PCR are decycling. The Mykkeltveit [15] and Champarnaud [2] sets are two examples of MDS constructions.

### 2.3 Sketching methods

The following sketching methods are commonly used by bioinformatics software packages and satisfy the sketching model described above.

The original *window-based* sketching methods are the minimizers [19, 20]. The input to the minimizers function is a context of *w k*-mers (originally called a window), and the function returns the index of the lexicographically smallest *k*-mer among the input, which is defined by some pre-determined order on the *k*-mers. Various refinements of minimizers exists such as local schemes [22], minmers [8]. These methods share the important property that they have a context or window length *>* 1 and they always select at least one *k*-mer in each input window. Consequently, these methods have a strong window guarantee of length equal to the input window length. In some applications, post-processing is applied to not select two identical homopolymer *k*-mers (say `AA` …`A`) in a row, to avoid selecting too many *k*-mers in low entropy regions of the genome. This effectively gives a relaxed window guarantee.

*Positional minimums* methods such as syncmers [4] have a context of length 1. They have extra parameters *s* and *t*, where a *k*-mer *m* is selected if the smallest *s*-mer among the *k−s* + 1 *s*-mers of *m* starts at position *t*. As before, the smallest *s*-mer is defined by some pre-determined order on *s*-mers. A notable generalization of syncmers is the parameterized syncmers [3] that use a bit mask of locations for the smallest *s*-mer instead of the parameter *t*.

When *s*-mers are tied, these methods use a left-most tie breaking rule. This rule has an interesting consequence on the window property. If *t* = 0 (i.e., the first base), then syncmers have a strong window guarantee, though the length might be as long as *σ*^*s*^ [13]. When 0 *< t ≤ s/*2 and the *s*-mer *m*_*s*_ at position *t* is minimal but not selected because an identical *s*-mer starts at position 0 *≤ i < t*, then *m*_*s*_ must be a “sesqui-power,” i.e., a word of the form *m*_*s*_ = *x*^*n*^*y* where *x* is of length *t− i* and *y* is a prefix of *x* (see Figure 1). Effectively this skips low-entropy genome regions that are repetitive with repeat lengths at most *t*, giving a relaxed window guarantee. When *t ≥ s*, there is no longer a window guarantee. Similarly, parameterized syncmers have a strong window guarantee if the bit 0 is set and either a relaxed guarantee or no guarantee for any other value of the mask.

**Figure 1:**
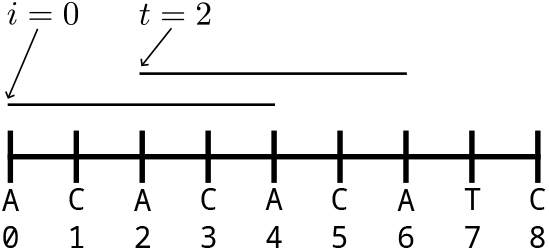
The left-most tie breaking rule implies a relaxed window guarantee for syncmers when t ≤ s/2. In this example k = 9, t = 2, s = 5, and the alphabetic order on 5-mers is used. The 9-mer is not selected because the 5-mer at position 2 is equal to the 5-mer at position 0 and the left-most tie breaking rule. Because this 5-mer overlaps with itself, it is “almost” repetitive: it is a sesqui-power x^n^y with x = `AC`, y = `A`, n = 2. Repetitive sequences of length 2 are skipped over.

*Hash-based* sketching methods are context-free functions that select a random subset of *k*-mers using a hash function. Examples include when the hash value is equal to 0 mod *p* or when it is less than or equal to *f* for a pre-determined modulus *p* or fraction *f*. In general, these sketching methods do not have any window guarantee.

## 3 Decycling in *k*-nonical space

In this section, unless specified otherwise, we assume that the sketching function *φ* is context-free.

### 3.1 Symmetric selection function

Compared to classic string algorithms, bioinformatics algorithms must take into account the double-stranded nature of DNA. In particular, when DNA is sequenced it is usually unknown which strands of the DNA are actually sequenced. In other words, the sequence of a read *r* and its reverse complement sequence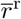carry the same information. Consequently, in algorithms using sketching, it is required to have “symmetric” selection functions, where the same decision is applied to a *k*-mer and its reverse complement. Formally, a selection function is defined to be symmetric when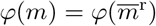for every *k*-mer *m*.

Sketching schemes are designed as classical string algorithms, ignoring the required symmetry. Then, in practice, software programs use canonical *k*-mers rather than *k*-mers. Effectively, instead of using the selection function *φ* they use the *canonicalized function φ*^c^ defined by *φ*^c^(*m*) = *φ*(*m*^c^). *φ*^c^ is guaranted to be symmetric.

It is often assumed that the canonicalized function *φ*^c^ essentially has the same properties as *φ*, and if anything, *φ*^c^ would select more *k*-mers than *φ*. In particular, a commonly-held belief is that, if the selection function *φ* has a (relaxed) window guarantee, then so too does *φ*^c^. Unfortunately this does not always hold, especially for the usual context-free methods. Even if a selection function *φ* has a strong window guarantee, the corresponding *φ*^c^ may not have any window guarantee.

### 3.2 Canonicalized decycling set

In the context-free case, a selection function is equivalent to a set of *k*-mers, and the canonicalized function is obtained via a simple set operation. Using this point of view, we clearly show in the next two Lemmas why

*φ*^c^ may not have the same decycling property as *φ*.

#### Lemma 1

(Canonicalized function). *The canonicalized selection function φ*^*c*^ *of φ is defined by the following set:*

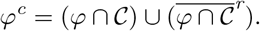

*Proof*. By definition of *φ*^c^, for any *k*-mer *m, m* ∈ *φ*^c^ ⟺ *m*^c^ ∈ *φ*. We consider two cases:

- when *m* ∈ 𝒞: *m* ∈ *φ*^c^ ⟺ *m* ∈ *φ*. That is, *φ*^c^ *∩ 𝒞* = *φ ∩ 𝒞*.
- when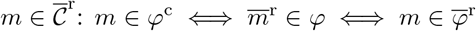. That is, 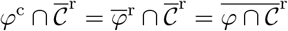.
- Because 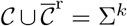 covers all the *k*-mers, the union of these two cases gives the desired result.

#### Lemma 2.

*Let* 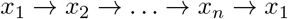*be a PCR. Then the reverse complemented k-mers form the PCR* 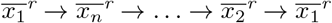. *These two PCRs may not be disjoint, in which case they are the same PCR. Proof*. The second part comes from the fact that PCRs partition the nodes of the de Bruijn graph.

For the first part, it is sufficient to show that for any *k*-mer *m* there is an edge 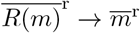where *R*(*m*) is the rotation of *m*. With *m* = *m*_0_*m*_1_ … *m*_*k−*1_, *R*(*m*) = *m*_1_ … *m*_*k−*1_*m*_0_, then 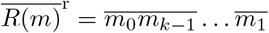 has an edge to 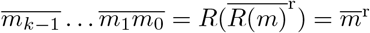.

Figure 2 shows examples of PCRs for the binary alphabet and *k* = 6. In this representation, the dashed vertical line acts as a “line of symmetry,” where canonical *k*-mers are on the left and non-canonical *k*-mers are on the right. Cases (a) and (c) are when the PCR and the reverse complement PCR are disjoint. Case (a) is when the PCR is entirely contained in 𝒞 or 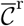, i.e., neither the PCR nor the reverse complement cross the dashed line. In case (c), both the PCR and the reverse complement cross the dashed line. Case (b) is when the same PCR contains *k*-mers and their reverse complements. *k*-mers on the dashed line are self-reverse complements.

**Figure 2:**
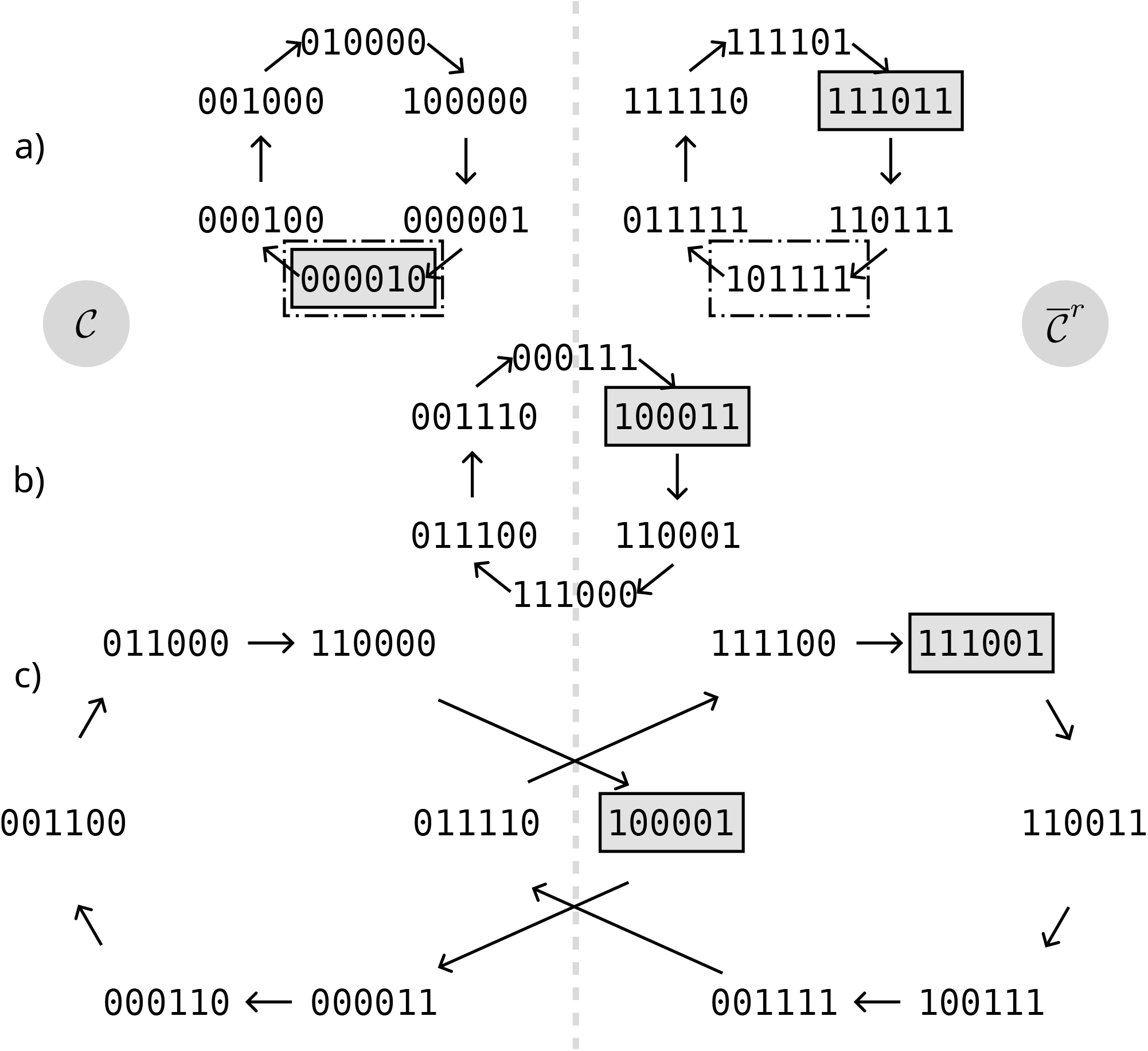
For k = 6 and ∑ = 2, examples of PCRs. Every k-mer on the left side is a canonical k-mer (∈ C), and k-mers symmetrical compared to the vertical line are reverse complement of each other. The k-mers inside gray boxes are an example of set φ and the k-mers in dashed boxes is the corresponding φ^c^ set. `000010` is both in φ and φ^c^. Not every PCR is covered by φ^c^ and consequently φ^c^ is not decycling.

The set *φ*^c^ is constructed from *φ* in two steps: (1) only the *k*-mers of *φ* on the left of the line are selected, and (2) the symmetrical *k*-mers are added. Consequently, in the cases where the *k*-mers of *φ* for a PCR and its reverse complement are in 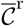 (on the right side of the plane), they are not selected in step 1 and *φ*^c^ does not cover these PCRs (see (b) and (c) in Figure 2). Hence *φ*^c^ does not necessarily contain a *k*-mer from every PCR, and it is therefore “less decycling” than the original function *φ*: the strongly connected components in SCC(*φ*^c^) are super-sets of SCC(*φ*).

Table 1 shows the number of strongly connected components and their cumulative size as a percent of the total number of *k*-mers (*k* = 15) for a variety of context-free schemes. The Mykkeltveit and Champarnaud sets are two known construction methods for decycling sets of minimum size. Although these sets are not used on their own as sketching methods, the Mykkeltveit set in particular has been used as a starting point to define sketching methods [17, 16, 5, 18]. By construction, these sets are decycling.

**Table 1:** For σ = 4 and k = 15, the table gives the number of SCCs and the percent of k-mers of the de Bruijn graph that are in a SCC. Syncmers use s = 6 and t ∈ {0, 1, 7}. Because Mykkeltveit, Champarnaud and Syncmers with t = 0 are decycling, there are no SCCs. Syncmers with t = 1 has a relaxed window guarantee that allows the homopolymer sequences, hence σ = 4 SCCs. In every case, the canonicalized set leaves one large strongly connected component in B_k_ ∖φ^c^.

|  | champarnaud |  | frac |  | mykkeltveit |  | syncmer 0 |  | syncmer 1 |  | syncmer 7 |  |
| --- | --- | --- | --- | --- | --- | --- | --- | --- | --- | --- | --- | --- |
| | $\varphi$ | $\varphi^c$ | $\varphi$ | $\varphi^c$ | $\varphi$ | $\varphi^c$ | $\varphi$ | $\varphi^c$ | $\varphi$ | $\varphi^c$ | $\varphi$ | $\varphi^c$ |
| Number of SCCs | 0 | 1 | 1 | 1 | 0 | 1 | 0 | 1 | 4 | 3 | 1337 | 1 |
| Cumulative size (%) | 0.00 | 95.94 | 89.98 | 89.98 | 0.00 | 84.35 | 0.00 | 87.23 | 0.00 | 84.19 | 0.01 | 83.74 |

The syncmer methods use *s* = 6 and values of *t ∈* {0, 1, 7 }, along with a random ordering of the *s*-mers. For *t* = 0 and *t* = 1 the syncmers have a strong and relaxed window guarantee (4 SCCs corresponding to the hompolymer sequences) respectively. For *t* = 7, because *t > s*, there is no window guarantee, and this set leaves many small strongly connected components. The value *t≈ k/*2 is an often recommended setting for syncmers [23].

The fractional set selects a fraction of *k*-mers *f* = 1*/*(*k−s* + 1) (i.e., it is expected to be equally as sparse as the syncmer methods) and uses a random permutation of *k*-mers. This method selects a random subset of all *k*-mers, has no window guarantee, and leaves one very large strongly connected component. It shows that a random set of *k*-mers does not approximate a set with a window guarantee.

After canonicalization, the window guarantees (whether strong or relaxed) do not hold anymore, and the sizes of the strongly connected components range from 83% to 95% of the de Bruijn graph. The Champarnaud set leads to a much larger component than the Mykkeltveit set, showing that the effect of canonicalization can vary significantly between methods, even with strong window guarantee. Canonicalization does not have a visible effect on the fractional set as it has no guarantee and a large SCC even before canonicalization.

### 3.3 Context-free vs. context-full methods

Although it is not a requirement of the definition of *φ*, every sketching methods with a context used in practice always selects at least one *k*-mer from their context (e.g., minimizers, minmers). As a consequence, for these sketching methods, the canonicalized function *φ*^c^ defined by 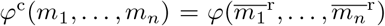 also always select at least one *k*-mer from its context. In other words, both *φ* and *φ*^c^ have, by construction, a strong window guarantee that is equal to the number of *k*-mers in the context.

The situation is very different with context-free methods: any non-trivial context-free sketching method must not always select a *k*-mer from the context. In fact, there is only one context-free method that always select the one *k*-mer from every context: it is the function that selects every *k*-mer from any input sequence and is equivalent to no sketching at all. Therefore, as seen in the previous section, a context-free method is not guaranteed to be a decycling set before or after canonicalization.

### 3.4 Canonicalized sketching deserts

The existence of large SCCs for a given sketching method implies that there exist long—even infinitely long—sequences without any selected *k*-mers. Moreover, the full set of sequences spelled-out by these SCCs is usually not known due to the complexity of the components. We call a sufficiently long region of a genome that does not contain any *k*-mers from a sketching method a *sketching deserts*. These regions are either undersampled or skipped by the sketching method. Thus, they are ignored by analyses using these sketching methods.

What precisely constitutes a sketching desert depends on the particular bioinformatics application and the type of data analyzed. Consider an application using *k*-mers from a sketch as anchors for alignment against a genome [10, 6] or using the sketches in an alignment-free method to characterize bacterial strands (e.g., Kraken [25]). In such applications, a minimum number of selected *k*-mers may be needed to (a) pass quality filters and (b) have significant statistical power. For example, with short-read sequencing (200 or fewer bases) and requiring a minimum of 4 selected *k*-mers per read, gaps of 50 bases or more between selected *k*-mers are problematic. Thus, regions of the genome of 50 bases or more without selected *k*-mers are considered sketching deserts for such an application.

Table 2 shows the cumulative size of the sketching deserts for two *k*-mer sizes in a few chromosomes (full table available in Appendix 8) of the human genome (GRCh38). For the *k* = 15 case, the size of sketching deserts can jump from a tiny fraction of a chromosome (0.02% for syncmer 0 and chromosome 1) to a non-negligeable fraction (1.16% with that same example) when switching to canonical *k*-mers. The potential size of sketching deserts increases with the length of the *k*-mers, as seen with *k* = 31 compared to *k* = 15.

**Table 2:** Cumulative size in mega-bases of the sketching desert of length ≥ 50 and ≥ 75 respectively for k = 15and k = 31 in the human genome GRCh38. For methods using a random order (fractional, syncmers), the valuesare the mean over 3 independent runs. The syncmer method parameters are k = 15, s = 6 and k = 31, s = 10 and t ∈ {0, 1, k/2}. The fractional parameter is set to match the number of selected k-mers as the syncmer method (f = 1/(k − s + 1)). In every method except fractional the canonicalized set has sketching desert orders of magnitude larger than the original method.

| | Chr | Mykkeltveit | | Champarnaud | | fractional | | syncmer 0 | | syncmer 1 | | syncmer $k/2$ | |
| --- | --- | --- | --- | --- | --- | --- | --- | --- | --- | --- | --- | --- | --- |
| | | $\varphi$ | $\varphi^c$ | $\varphi$ | $\varphi^c$ | $\varphi$ | $\varphi^c$ | $\varphi$ | $\varphi^c$ | $\varphi$ | $\varphi^c$ | $\varphi$ | $\varphi^c$ |
| $k = 15, \geq 50$ | 1 | 0.02 | 1.28 | 0.07 | 79.02 | 8.07 | 8.16 | 0.04 | 2.88 | 0.01 | 2.17 | 0.16 | 2.06 |
|  | 10 | 0.01 | 0.74 | 0.04 | 45.82 | 4.63 | 4.73 | 0.03 | 1.69 | 0.01 | 1.24 | 0.09 | 1.17 |
|  | 22 | 0.00 | 0.20 | 0.01 | 13.01 | 1.40 | 1.39 | 0.01 | 0.48 | 0.00 | 0.38 | 0.03 | 0.33 |
|  | X | 0.01 | 0.83 | 0.05 | 52.70 | 5.41 | 5.53 | 0.03 | 1.89 | 0.01 | 1.47 | 0.12 | 1.37 |
| $k = 31, \geq 75$ | 1 | 1.98 | 10.82 | 3.00 | 103.06 | 63.20 | 38.14 | 3.49 | 18.74 | 2.34 | 17.68 | 0.29 | 13.65 |
|  | 10 | 1.15 | 6.23 | 1.71 | 59.28 | 24.68 | 57.52 | 2.02 | 11.19 | 1.38 | 10.05 | 0.17 | 8.02 |
|  | 22 | 0.33 | 1.82 | 0.52 | 16.69 | 8.90 | 8.37 | 0.56 | 3.26 | 0.40 | 2.93 | 0.05 | 2.39 |
|  | X | 1.48 | 7.32 | 1.90 | 70.24 | 95.06 | 62.54 | 2.39 | 12.54 | 1.58 | 11.88 | 0.21 | 8.93 |

The actual length of these sketching desert results from the particular interaction between the genome sequence and the sequence spelled by the strongly connected components of the method. The canonicalized Champarnaud set for example has a bad interaction with the human genome sequence and has very large sketching deserts. Although this set probably has not been used in practice, it shows the dramatic effect canonicalization can have. The effect on the syncmer methods for these random choices of orders on the *s*-mers is less dramatic, although still quite large, especially for large *k*-mers. On the other hand, the relationship between the order used on *s*-mers and the size of the sketching desert is completely unknown and nothing in the method a priori prevents similarly large effects when using an order on *s*-mers that interacts poorly with the human genome.

The fractional set method has the weakest guarantee, with significant sketching deserts before and after canonicalization. The variance due to the choice of the random permutation on *k*-mers is also high with this method (standard deviation of 34.7 Mb for *φ* and 28.3 Mb for *φ*^c^ on Chr 1 with *k* = 31).

Similar effects are observed on the sequence of protein-coding genes. For example, with syncmers (*k* = 15, *s* = 6, *t* = 0), one of the random orders on *s*-mers leaves *>* 20% of the sequence of MIER2 (transcript MIER2-201 with 7084 bases) in sketching deserts, with multiple gaps of *>* 100 bases, when using canonicalization. Without canonicalization and the same order there is no gap of 50 bases or more, hence no sketching deserts. Even more pronounced effect are observed with the longer *k*-mers: gene RBBP9 (transcript RBBP9-203 with 1121 bases) has *>* 90% of its sequence in sketching deserts for syncmers (*k* = 31, *s* = 10, *t* = 1) in *k*-nonical space while having no desert before canonicalization.

This also illustrates the impact of the random choice of the order on *s*-mers. For the same syncmers parameters, while for all 3 tested orders there exist transcripts with sketching deserts of *>* 20% of their length, the MIER2 gene has sketching deserts for only one of these orders.

### 3.5 The union set and sparse canonicalization

In the formula of Lemma 1, the intersection with the set of canonical *k*-mers 𝒞 is the reason why *φ*^c^ intersects with fewer cycles of the de Bruijn graph, creating larger strongly connected components and larger sketching deserts. By avoiding this initial intersection, the following simple method creates a symmetric selection function from a given selection function *φ*:

#### Definition 1

(Union function). *As a set, the “union” selection function is defined by:*

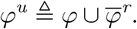

That is a *k*-mer *m* is selected by *φ*^u^ if either *m* or its reverse complement 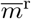 is selected by *φ*. Because *φ*^u^ is a super-set of *φ*, using *φ*^u^ does not introduce any new strongly connected components or sketching deserts. This union function is not entirely new. Although motivated by a different goal, it is suggested in [18] to use an order for minimizers which is based on the set *φ*^u^, where *φ* is the Mykkeltveit set.

On the other hand, the union selection function is not traditionally used with sketching methods, because the size of the set *φ*^u^ is likely about double that of *φ*, which could change the behavior of the sketching method. For example, the number of selected *k*-mers in a sequence *S*—approximately |*φ*^u^∩ 𝒦(*S*) | —is likely to double as well, reducing the sparsity of sketches created by the union sketching function compared to that of the original function (i.e., it affects the *density* [12, 11] of the sketching method). By comparison, the canonicalized function, thanks to the intersection with 𝒞 which contains slightly over half of all the *k*-mers, is expected to have a size similar to that of the original sketching function.

We therefore propose the following optimization problem of finding a function which is as close as possible to the canonicalization function while not increasing the number of strongly connected components.

#### Problem 1

(Sparse canonicalization). *Given a set φ, find the smallest set A such that φ*^*c*^ *⊂ A and* SCC(*A*) *⊂* SCC(*φ*).

The union set (Definition 1 satisfies the two conditions of Problem 1 but it is not necessarily of smallest size. We use the following greedy procedure to reduce its size: starting from a union set *φ*^u^, each *k*-mer from *φ*^u^ is tested and removed if its removeal does not break the decycling property. For example, with *σ* = 4, *k* = 6 the Mykkeltveit set contains 700 *k*-mers and the greedy procedure removes 123 *k*-mers from the associated union set while still being decycling. Performance of this procedure is shown in Section 4.3.

## 4 Symmetric sketching function design

The previous section focused on the effect of using canonical *k*-mers with an existing sketching method. An alternative approach is to directly design a sketching method in “*k*-nonical space,” i.e., finding selection functions *φ* that satisfy 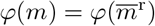 by design. In this section, we develop an efficient ILP procedure that finds a set of *k*-mers of smallest size under the constraints that it satisfies the symmetry condition and has a strong window guarantee of the desired length (i.e., it is decycling and the longest path in the DAG B _*∖*_*φ* is less than some specified parameter).

### 4.1 Feedback Vertex Sets and Nilpotency

A feedback vertex set (FVS) of a digraph *G* is a set of vertices *W ⊆ V* such that the digraph *G ∖ W* induced by the deletion of *W* is a directed-acyclic graph. The Minimum Feedback Vertex Set (MFVS) problem is to find the minimum cardinality FVS and was one of the original 21 problems shown to be NP-hard [7].

Every MDS solves the MFVS problem on the de Bruijn graph, yet unlike the general problem, there are efficient algorithms to find MDSs (i.e., the Mykkeltveit [15] and Champarnaud [2] algorithms). However, the space of MDSs is empirically observed to offer substantial diversity over optimization metrics like the maximum remaining path length, with efficiently selected sets falling short of global optima [13]. Moreover, there are no known algorithms to create symmetric minimum size decycling sets, i.e., sets *M* such that 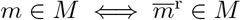.

To optimize MDSs over both the standard and canonical de Bruijn graph, we develop an integer linear programming (ILP) formulation for a variant of the MFVS problem that (a) accepts a maximum remaining path length constraint, (b) works with symmetries such as the reverse complement, and (c) minimizes the expected remaining path length after decycling.

Our formulation uses a connection between DAGs and nilpotent matrices to enforce a maximum path length and compute the expected path length. A matrix *A∈* ℝ^*n×n*^ is *nilpotent* when *A*^*t*^ = 0 for some *t ∈* ℕ. The minimal *t* is called the *nilpotent index* of the matrix and is always less than or equal to *n* when it exists [1]. When *A* is nonnegative (i.e., it has no negative entries), there is a convenient equivalent condition to nilpotence defined with the vector *e* of all 1 ‘s.

#### Lemma 3.

*A nonnegative matrix A is nilpotent if and only if A*^*t*^*e* = 0 *for some t≤ n. The smallest such t is the nilpotent index of A*.

*Proof*. If *A* is nilpotent, we have for some *t ≤ n* that *A*^*t*^*e* = 0(*e*) = 0. If *A* is nonnegative and *A*^*t*^*e* = 0, it follows that *A*^*t*^ is the zero matrix because (1) *A*^*t*^*e* is the vector of row sums, (2) a nonnegative vector sums to zero if and only if it is the zero vector, and (3) nonnegative matrices are closed under matrix multiplication. In both directions, *t* can be taken to be minimal with no change, so the smallest such *t* is precisely the nilpotent index.

Suppose *G* = (*V, E*) is a digraph with *V* = {1, 2, …, *n*}. Let *A ∈* ℝ^*n×n*^ be the adjacency matrix of *G*, defined as *A*_*ij*_ = 1 if (*i, j*) *∈ E* and 0 otherwise. A walk in *G* is a string of vertices *v*_0_*v*_1_ … *v*_*t*_ such that consecutive vertices are adjacent, i.e., (*v*_*i*_, *v*_*i*+1_) *∈ E*. A path is a walk with no repeated vertices. Recall that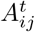 is the number of distinct walks of length *t* between nodes *i* and *j*.

#### Lemma 4.

*G is a DAG if and only if A is nilpotent. Moreover, when G is a DAG, the nilpotent index is one more than the longest path length*.

*Proof*. Suppose *G* is a DAG. Every walk in *G* is a path because a walk with a repeated vertex would imply the existence of a cycle. As the vertex set is finite, there is a longest path of length *t*^*\**^ *< n*. Thus,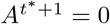 since a non-zero entry would indicate a path longer than the longest path. This is the nilpotent index because there are sub-paths of the longest path for every length less than or equal to *t*^*\**^.

Now suppose *G* has a cycle. Then *G* also has a walk of every length constructed by starting at any vertex in the cycle and traversing it until reaching the desired length. Thus, *A*^*t*^*≠*0 for any *t* because at least one entry must be greater than zero to count the walks around the cycle, showing that *A* is not nilpotent.

The *maximum t-nilpotent submatrix* (*t*-MNS or MNS) problem is to find a nilpotent submatrix of *A* that maximizes selection with respect to a weight vector *w*, while ensuring the submatrix has a nilpotent index no larger than *t*. The general version of this problem has not been studied, but it is equivalent to MFVS when

*A* is an adjacency matrix and *t* = *n*. Moreover, the nonnegative case of *t*-MNS is equivalent to MFVS with an additional max path-length constraint for values of *t < n*.

#### Lemma 5.

*MNS is NP-hard via a reduction from MFVS*.

*Proof*. Let *G* be a digraph represented by its adjacency matrix *A*, and let Γ *⊆* [*n*] be the indices of a nilpotent submatrix of *A*. By Lemma 4, the induced subgraph *G*_Γ_ is a DAG, so the complement [*n*] *−* Γ is a FVS. The same argument taken in reverse shows that a FVS corresponds to a nilpotent submatrix. Thus, every FVS is the complement of an index set for a nilpotent submatrix of *A*. This shows that solutions to MNS for *A* are equivalent to solutions to MFVS for *G* up to a complement.

### 4.2 MDS-selection ILP

From the proof of Lemma 5, we see that finding a minimum FVS is equivalent to solving *n*-MNS with an adjacency matrix. Moreover, if we constrain the nilpotent index, we constrain the maximum remaining path length of the selected vertices. Thus, we now develop a formulation for the MNS problem over nonnegative matrices that we will use to find path-constrained MDSs. (See Corollary 1 in Appendix 9 for an extension of our formulation to the general case.)

For the following, *A* is an *n × n* matrix, Γ is an index set, *A*_Γ_ is the submatrix with rows and columns indexed by Γ, *e*_Γ_ is the vector of all 1s indexed by Γ, and [*n*] is the set of indices from 1 to *n*.

#### Theorem 1.

*The following formulation solves for a* (*t* + 1)*-nilpotent submatrix, maximizing selection with respect to a weight vector w:*

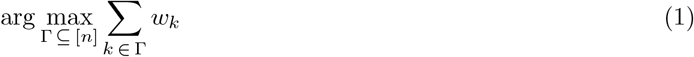

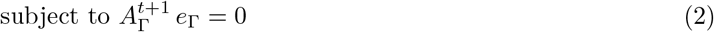

*Moreover, it can be formulated as a mixed-integer linear program with 𝒪* (*n*) *binary variables, 𝒪* (*t·n*) *continuous variables, and 𝒪* (*t · n*) *constraints*.

*Proof*. By Lemma 3 ‘s result, the constraint ensures the feasible set consists precisely of all nilpotent submatrices with nilpotent index *t* + 1 or smaller. Thus, solving this problem is equivalent to finding the nilpotent submatrix maximizing selection with respect to *w*.

We can use a binary vector of size *n* to represent Γ. In Appendix 9 and Figure 3 we describe how to construct a (*t* + 1)-layer circuit, where layer *j* represents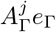. Each layer requires 𝒪(*n*) continuous variables and 𝒪 (*n*) constraints. Thus, we have 𝒪(*n*) binary variables, 𝒪(*t · n*) continuous variables, and (*t · n*) constraints, as claimed.

**Figure 3:**
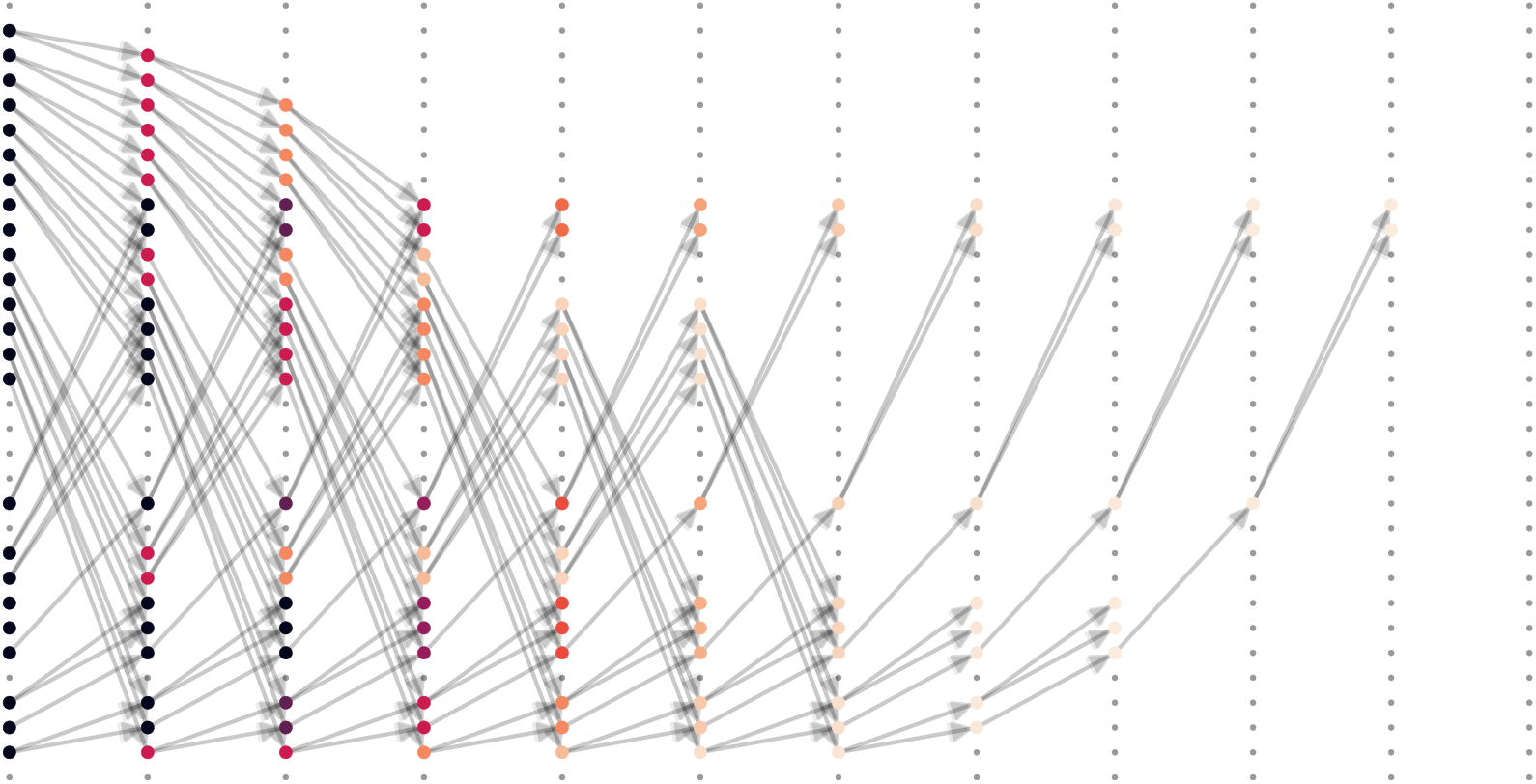
Shown is a solution found by the ILP for the σ = 2 and k = 5 de Bruijn graph. In this representation, every column of dots represent a copy of the de Bruijn graph with σ^k^ dots per column each representing a k-mer. There is as many columns as the upper-bound on the remaining path length. Dots have at most σ out-going edges and nodes with in- and out-degree 0 are the selected nodes of the set. There is an edge between two nodes from a column to the next iff neither node is selected in the set. The selected nodes represent a MDS that also (a) minimizes the remaining max path length (11) and (b) among those minimizes the expected path length. The ILP variable associated with the dot at row i column j is the probability that a path of length j ends at vertex i of B_k_. Vertices are colored such that the small dots have probability zero, lighter nodes have a lower probability, and darker nodes have a higher probability.

When *A* is an adjacency matrix, this formulation solves MFVS with respect to the weights given by *w*. With an appropriate modification to *A*, this framework also models the expected path length of a uniform random walk that stops once a removed vertex is hit.

To do this, we use the random-walk adjacency matrix defined as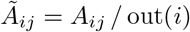, where out(*i*) is the number edges originating at vertex *i*. The entry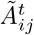 is the probability that a random walk starting at vertex *i* is at vertex *j* during time *t* when uniformly sampling an outgoing edge. For an index set Γ,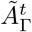 is also the probability that a random walk starting at vertex *i* is at vertex *j* during time *t*, with the caveat that some states are absorbing, i.e., a random walk stops once a state in [*n*] *−* Γ is hit. This leads to two expectation identities given by the random walk.

For random-walk matrix *Ã*, index set Γ⊆ [*n*], and probability vector *p* ∈ ℝ^*n*^, the expected time before a random walk *X* is absorbed by Γ satisfies

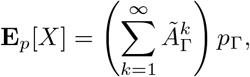

which is equal to infinity when the subset Γ does not asymptotically halt random walks (i.e., it does not represent a decycling set). When Γ induces a submatrix of nilpotent index *t* + 1 or smaller (i.e., it represents a decycling set), this simplifies to

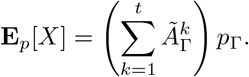

Thus, using the construction of Appendix 9, we can represent all the vectors used in these expectation formulas. We combine this with Theorem 1 to find a MFVS that secondarily minimizes the expected hitting time of a uniformly random walk using a rescaled uniform probability vector. In Figure 3, we visualize this process as a probabilistic circuit over the de Bruijn graph with *σ* = 2 and *k* = 5.

To work with symmetries, we use a change of basis that considers all equivalent vertices to be the same. For example, if node *v* is equivalent to *w* (e.g.,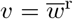), we simply replace both respective vectors with a new basis vector (*v* + *w*)*/*2 in the adjacency matrix construction.

### 4.3 Results

We tested our ILP formulation against our greedy union algorithm with the *σ* = 2 de Bruijn graph with *k* ranging from 5 to 8 for these experiment. This range of *k* was chosen because the ILP solved these instances (in order of *k*) in 2, 10, 30, and 58 minutes and required no more than one gigabyte of working memory. The ILP did not solve the *k* = 9 instance after 3 hours and peaked at 5 gigabytes of working memory. Because of the exponential growth of the problem size, further optimizations or algorithmic insights are necessary to be able to tackle the larger instances necessary for practical uses. E.g., for *σ* = 4 and *k* = 15 the de Bruijn graph has over a billion nodes and is out of reach for the current ILP.

Table 3 shows the results of our experiment. We see that the ILP optimized selections have, as expected, a smaller cardinality than the greedy selections. The maximum path lengths of the greedy optimizers are occasionally shorter because more *k*-mers were selected by greedy and the ILP optimizes first for the cardinality of the set.

**Table 3:** Statistics on the solution of the greedy procedure of Section 3.5 and the ILP method. Results are for the de Bruijn graph with σ = 2 and k ranging from 5 to 8. The columns show the cardinality of the selected sets, maximum path length B_k_ after removing the sets, and the rounded expected path length after removing the sets. The Mykkeltveit (labeled Mykk.) and Champarnaud (labeled Champ.) sets are used as the starting points of the greedy procedure. The ILP procedure minimizes (in order) for cardinality, maximum path length, and expected path length.

| k | Cardinality |  |  | Maximum Path Length |  |  | Expected Path Length |  |  |
| --- | --- | --- | --- | --- | --- | --- | --- | --- | --- |
|  | Champ. | Mykk. | ILP | Champ. | Mykk. | ILP | Champ. | Mykk. | ILP |
| 5 | 12 | 12 | 12 | 5 | 5 | 6 | 1.16 | 1.17 | 1.36 |
| 6 | 19 | 19 | 16 | 14 | 14 | 9 | 1.66 | 1.72 | 1.53 |
| 7 | 30 | 30 | 28 | 15 | 15 | 20 | 2.12 | 2.08 | 1.62 |
| 8 | 45 | 51 | 44 | 34 | 18 | 23 | 2.97 | 2.45 | 2.10 |

## 5 Conclusion

The use of canonical *k*-mers is the standard modification that allows sketching methods for standard text to work with biological sequences. We have shown this has a previously unrecognized flaw when applied to context-free sketching methods. This approach creates sketching deserts that make some sequences effectively invisible to downstream algorithms that use the sketch, potentially creating biases in the analysis. We described the theoretical mechanism behind the creation of these sketching deserts and provided two different options to designing sketching methods that properly handle sequences that are equivalent to their reverse complements. The first method modifies existing sketching methods, but unlike canonical *k*-mers, this modification does not create sketching deserts. The second designs *de novo* symmetric sketching methods that outperform their symmetrized counterparts.

In general, sketching methods that are used with genomics data must be explicitely designed and validated to handle the equivalence between a sequence and its reverse complement. This must be an intentional step in the design process, not an afterthought.

## 7 Acknowledgments

This work was supported in part by the US National Science Foundation [DBI-1937540, III-2232121], the US National Institutes of Health [R01HG012470] and by the generosity of Eric and Wendy Schmidt by recommendation of the Schmidt Futures program.

