## Appendix for "*k*-nonical space: sketching with reverse complements"

**Table 4:** *Sketching desert cumulative sizes for $\sigma = 4, k = 15$ and desert 50 bases or longer.*

| Chr | Mykkeltveit | | Champarnaud | | fractional | | syncmer 0 | | syncmer 1 | | syncmer 7 | |
|---|---|---|---|---|---|---|---|---|---|---|---|---|
| | $\varphi$ | $\varphi^c$ | $\varphi$ | $\varphi^c$ | $\varphi$ | $\varphi^c$ | $\varphi$ | $\varphi^c$ | $\varphi$ | $\varphi^c$ | $\varphi$ | $\varphi^c$ |
| 1 | 0.02 | 1.28 | 0.07 | 79.02 | 8.07 | 8.16 | 0.04 | 2.88 | 0.01 | 2.17 | 0.16 | 2.06 |
| 2 | 0.02 | 1.33 | 0.07 | 83.29 | 8.40 | 8.50 | 0.05 | 2.96 | 0.01 | 2.20 | 0.16 | 2.10 |
| 3 | 0.02 | 1.08 | 0.05 | 68.75 | 6.88 | 7.00 | 0.04 | 2.44 | 0.01 | 1.83 | 0.13 | 1.73 |
| 4 | 0.02 | 1.03 | 0.05 | 66.33 | 6.61 | 6.66 | 0.03 | 2.36 | 0.01 | 1.78 | 0.12 | 1.67 |
| 5 | 0.02 | 0.99 | 0.05 | 62.88 | 6.25 | 6.35 | 0.03 | 2.27 | 0.01 | 1.75 | 0.11 | 1.63 |
| 6 | 0.02 | 0.95 | 0.05 | 58.98 | 5.94 | 6.02 | 0.03 | 2.10 | 0.01 | 1.57 | 0.11 | 1.50 |
| 7 | 0.01 | 0.87 | 0.04 | 54.35 | 5.54 | 5.60 | 0.03 | 2.00 | 0.01 | 1.63 | 0.11 | 1.37 |
| 8 | 0.01 | 0.79 | 0.04 | 50.18 | 5.02 | 5.12 | 0.03 | 1.78 | 0.01 | 1.36 | 0.10 | 1.27 |
| 9 | 0.01 | 0.66 | 0.03 | 41.66 | 4.28 | 4.26 | 0.02 | 1.48 | 0.01 | 1.12 | 0.07 | 1.07 |
| 10 | 0.01 | 0.74 | 0.04 | 45.82 | 4.63 | 4.73 | 0.03 | 1.69 | 0.01 | 1.24 | 0.09 | 1.17 |
| 11 | 0.01 | 0.72 | 0.04 | 45.48 | 4.75 | 4.74 | 0.03 | 1.63 | 0.01 | 1.36 | 0.09 | 1.16 |
| 12 | 0.01 | 0.74 | 0.04 | 45.99 | 4.62 | 4.79 | 0.03 | 1.65 | 0.01 | 1.25 | 0.09 | 1.15 |
| 13 | 0.01 | 0.54 | 0.03 | 34.47 | 3.45 | 3.44 | 0.02 | 1.23 | 0.00 | 0.92 | 0.06 | 0.85 |
| 14 | 0.01 | 0.48 | 0.03 | 31.19 | 3.19 | 3.21 | 0.02 | 1.12 | 0.00 | 0.84 | 0.06 | 0.78 |
| 15 | 0.01 | 0.46 | 0.03 | 28.92 | 2.98 | 3.02 | 0.02 | 1.02 | 0.00 | 0.82 | 0.05 | 0.72 |
| 16 | 0.01 | 0.45 | 0.02 | 27.58 | 2.86 | 2.89 | 0.01 | 1.03 | 0.01 | 0.83 | 0.06 | 0.75 |
| 17 | 0.01 | 0.45 | 0.02 | 27.88 | 2.98 | 2.99 | 0.02 | 1.05 | 0.00 | 0.96 | 0.06 | 0.70 |
| 18 | 0.01 | 0.42 | 0.02 | 27.71 | 2.84 | 2.78 | 0.01 | 0.99 | 0.00 | 0.82 | 0.05 | 0.74 |
| 19 | 0.00 | 0.30 | 0.02 | 19.27 | 2.03 | 2.08 | 0.01 | 0.76 | 0.00 | 0.59 | 0.05 | 0.54 |
| 20 | 0.01 | 0.34 | 0.01 | 21.68 | 2.28 | 2.23 | 0.01 | 0.78 | 0.00 | 0.63 | 0.05 | 0.56 |
| 21 | 0.00 | 0.22 | 0.01 | 13.90 | 1.42 | 1.41 | 0.01 | 0.50 | 0.00 | 0.39 | 0.03 | 0.35 |
| 22 | 0.00 | 0.20 | 0.01 | 13.01 | 1.40 | 1.39 | 0.01 | 0.48 | 0.00 | 0.38 | 0.03 | 0.33 |
| X | 0.01 | 0.83 | 0.05 | 52.70 | 5.41 | 5.53 | 0.03 | 1.89 | 0.01 | 1.47 | 0.12 | 1.37 |
| Y | 0.00 | 0.13 | 0.01 | 8.19 | 0.86 | 0.86 | 0.00 | 0.32 | 0.00 | 0.22 | 0.03 | 0.24 |

### Appendix

#### 8 Complete sketching desert tables

Table 4 and 5 show the sizes of the sketching deserts for all the chromosomes of the human reference genome GRCh38.

#### 9 ILP Circuit

##### 9.1 Intuition

A *Krylov sequence* of order $t$ for matrix $A$ and vector $x$ is the sequence $(x, Ax, \ldots, A^t x)$. Such sequences frequently arise in numerical linear algebra to optimize vector selection and capture cyclic dynamics [24]. For example, when $x$ is a probability vector and $A$ is a random-walk matrix, Krylov sequences describe the distribution of random walk states as they evolve over time.

In our case, we work with random walk matrices that have blocking or absorbing states. In this case, the Krylov sequence describes the location of escaping walks over each time step. We interpret this as a probabilistic circuit, where each column contains a copy of all vertices and each row represents the time evolution of that vertex. For an order $t+1$ circuit, the final column consists entirely of zeros if and only if a FVS with a maximum remaining path length of $t$ or less is selected. We show an example of this for the de Bruijn with $\sigma = 2$ and $k = 12$ in Figure 4.

##### 9.2 Circuit Construction

We now show how to construct an ILP representation of a Krylov sequence for a fixed nonnegative matrix $A$ and a fixed nonnegative vector $x$ where we are given binary variables that represent the submatrix. For the

**Table 5:** *Sketching desert cumulative sizes for $\sigma = 4, k = 31$ and desert 75 bases or longer.*

| Chr | Mykkeltveit | | Champarnaud | | fractional | | syncmer 0 | | syncmer 1 | | syncmer 15 | |
|---|---|---|---|---|---|---|---|---|---|---|---|---|
| | $\varphi$ | $\varphi^c$ | $\varphi$ | $\varphi^c$ | $\varphi$ | $\varphi^c$ | $\varphi$ | $\varphi^c$ | $\varphi$ | $\varphi^c$ | $\varphi$ | $\varphi^c$ |
| 1 | 1.98 | 10.82 | 3.00 | 103.06 | 63.20 | 38.14 | 3.49 | 18.74 | 2.34 | 17.68 | 0.29 | 13.65 |
| 2 | 2.16 | 11.43 | 3.06 | 108.94 | 81.30 | 34.32 | 3.66 | 19.89 | 2.52 | 18.34 | 0.30 | 14.16 |
| 3 | 1.72 | 9.44 | 2.47 | 90.30 | 44.12 | 20.19 | 2.95 | 16.71 | 2.06 | 15.27 | 0.24 | 11.78 |
| 4 | 1.70 | 9.13 | 2.35 | 87.60 | 53.67 | 81.03 | 2.84 | 15.60 | 1.97 | 14.39 | 0.24 | 11.22 |
| 5 | 1.62 | 8.62 | 2.21 | 82.69 | 44.36 | 43.77 | 2.83 | 15.15 | 1.90 | 13.75 | 0.23 | 11.12 |
| 6 | 1.49 | 8.11 | 2.16 | 77.65 | 31.59 | 24.38 | 2.54 | 14.18 | 1.74 | 12.99 | 0.20 | 10.13 |
| 7 | 1.39 | 7.36 | 2.05 | 71.54 | 42.75 | 20.41 | 2.34 | 13.31 | 1.58 | 12.21 | 0.19 | 9.19 |
| 8 | 1.28 | 7.01 | 1.83 | 65.66 | 33.71 | 19.37 | 2.16 | 12.13 | 1.52 | 11.21 | 0.18 | 8.57 |
| 9 | 1.06 | 5.93 | 1.54 | 54.98 | 22.34 | 14.81 | 1.81 | 10.11 | 1.23 | 9.09 | 0.14 | 7.19 |
| 10 | 1.15 | 6.23 | 1.71 | 59.28 | 24.68 | 57.52 | 2.02 | 11.19 | 1.38 | 10.05 | 0.17 | 8.02 |
| 11 | 1.16 | 6.56 | 1.71 | 59.61 | 27.13 | 16.91 | 2.02 | 11.21 | 1.35 | 10.50 | 0.16 | 8.08 |
| 12 | 1.18 | 6.25 | 1.73 | 59.96 | 27.06 | 15.82 | 1.97 | 11.07 | 1.34 | 10.38 | 0.19 | 7.76 |
| 13 | 0.88 | 4.74 | 1.21 | 45.24 | 25.19 | 36.26 | 1.47 | 8.08 | 1.00 | 7.57 | 0.12 | 5.87 |
| 14 | 0.79 | 4.31 | 1.15 | 40.94 | 13.26 | 40.35 | 1.37 | 7.67 | 0.94 | 6.82 | 0.11 | 5.27 |
| 15 | 0.79 | 3.97 | 1.10 | 37.61 | 18.03 | 12.17 | 1.24 | 6.90 | 0.86 | 6.42 | 0.10 | 4.98 |
| 16 | 0.71 | 3.84 | 1.10 | 35.66 | 23.52 | 8.23 | 1.19 | 6.74 | 0.85 | 6.06 | 0.10 | 5.02 |
| 17 | 0.95 | 4.01 | 1.14 | 35.14 | 20.66 | 36.59 | 1.19 | 6.92 | 0.88 | 6.21 | 0.10 | 4.80 |
| 18 | 0.68 | 3.82 | 0.95 | 36.21 | 11.38 | 51.20 | 1.33 | 6.58 | 0.78 | 6.09 | 0.09 | 4.73 |
| 19 | 0.48 | 2.54 | 0.85 | 24.23 | 12.52 | 20.96 | 0.85 | 4.85 | 0.62 | 4.47 | 0.08 | 3.52 |
| 20 | 0.58 | 3.24 | 0.81 | 28.17 | 13.04 | 8.70 | 0.93 | 5.26 | 0.68 | 4.79 | 0.08 | 3.94 |
| 21 | 0.36 | 1.92 | 0.49 | 18.14 | 6.86 | 5.47 | 0.59 | 3.35 | 0.42 | 3.04 | 0.05 | 2.44 |
| 22 | 0.33 | 1.82 | 0.52 | 16.69 | 8.90 | 8.37 | 0.56 | 3.26 | 0.40 | 2.93 | 0.05 | 2.39 |
| X | 1.48 | 7.32 | 1.90 | 70.24 | 95.06 | 62.54 | 2.39 | 12.54 | 1.58 | 11.88 | 0.21 | 8.93 |
| Y | 0.21 | 1.12 | 0.29 | 10.79 | 4.00 | 2.13 | 0.35 | 1.92 | 0.25 | 1.80 | 0.04 | 1.44 |

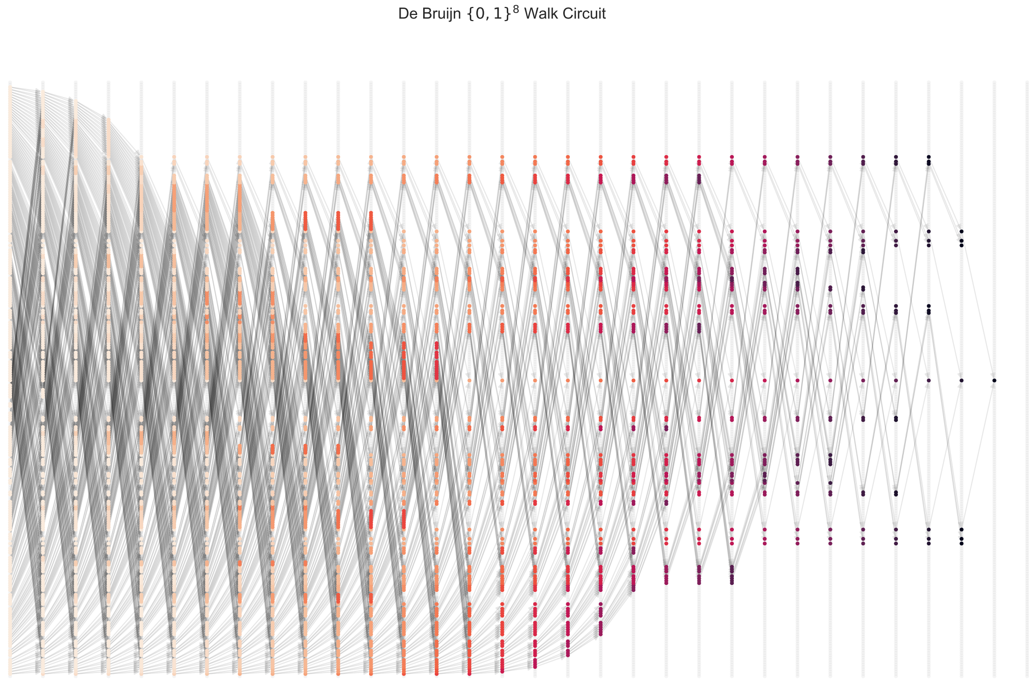


**Figure 4:** *Shown is the probabalistic decycling circuit for a $\sigma = 2$ and $k = 8$ de Bruijn graph MDS that (a) minimizes the remaining max path length (31) and (b) chooses among those minimizers one with a minimal expected path length. Vertices are colored such that translucent nodes are not occupied, darker nodes have a lower probability, and lighter nodes have a higher probability. Each column is an independent copy of the vertices, and each row shows how random walks occupy that vertex over time.*

following, $e$ is the vector of all 1's, $\|\cdot\|_\infty$ refers to the $\ell_\infty$ norm, $\neg\Gamma$ is the negation of the binary vector $\Gamma$, and $\odot$ is the Schur product, i.e., entrywise multiplication between vectors.

**Theorem 2.** *For a binary indicator vector $\Gamma \in \{0,1\}^n$, a nonnegative matrix $A \in \mathbb{Q}^{n\times n}$, and a fixed vector $v \in \mathbb{Q}^n$ such that $v_\Gamma = 0$, the vectors $\{A_\Gamma v_\Gamma, A_\Gamma^2 v_\Gamma, \ldots, A_\Gamma^t v_\Gamma\}$ are linearly representable with $\mathcal{O}(t \times n)$ continuous variables and $\mathcal{O}(t \times n)$ linear constraints.*

*Proof.* We show this via induction on $t$. We have entrywise bounds on the increase in magnitude given by $Av \le \|v\|_\infty Ae$. This holds because $v \le \|v\|_\infty e$ and multiplication by a nonnegative matrix preserves entrywise order. Representing this upper bound by $M$, the following constraints suffice to enforce that a nonnegative vector $x \in \mathbb{Q}_+^n$ satisfies $x_\Gamma = A_\Gamma v_\Gamma$ and $x_{\neg\Gamma} = 0$:

$$Av - M \odot \neg\Gamma \;\le\; x \;\le\; Av \;\text{ and }\; x \le M\Gamma.$$

We see from the hypothesis on $v$ that

$$(Av)_k = \sum_{j=1}^{n} A_{kj} v_j = \sum_{\Gamma_j=1}^{n} A_{kj} v_j = (A_\Gamma v_\Gamma)_k.$$

Thus, it follows that the constraints are both consistent and sufficient. To see this, suppose first that $\Gamma_k = 0$. Then

$$(Av)_k - M \;\le\; 0 \;\le\; x_k \;\le\; M\Gamma_k = 0 \le (Av)_k \implies x_k = 0.$$

On the other hand, when $\Gamma_k = 1$, we have

$$0 \le (Av)_k - M \le x_k \le (A_\Gamma v_\Gamma)_k \le M \implies x_k = (A_\Gamma v_\Gamma)_k.$$

This shows the construction is correct for $t = 1$. Moreover, we used $n$ variables and $3n$ constraints, which falls within the stated bounds.

Now suppose this is correct for up to $t-1$ steps. Then we have a variable $x$ such that $x_\Gamma = A_\Gamma^{t-1} v_\Gamma$ and $x_{\neg\Gamma} = 0$. Repeat the $t = 1$ construction with $x$ used as the vector, $A$ as the matrix, and a new variable vector $y$ as output. This is possible because $x_{\neg\Gamma} = 0$ by design and terms in the $t = 1$ constraints do multiplicatively depend on $x$, aside from $M$ which can be taken as $M_t = \|A\|_\infty^t \|v\|_\infty$ instead because $|(A^t v)_k| \le \|A^t v\|_\infty \le \|A^{t-1}\|_\infty \|A\|_\infty \|v\|_\infty \le \|A\|_\infty^t \|v\|_\infty$ This bound and the construction imply the new vector satisfies $y_\Gamma = A_\Gamma x_\Gamma = A_\Gamma^t v$ and $y_{\neg\Gamma} = 0$. Moreover, we add only $\mathcal{O}(n)$ new variables and $\mathcal{O}(n)$ new constraints.

In total, there are $(t-1)\,\mathcal{O}(n) + \mathcal{O}(n) = t\,\mathcal{O}(n)$ total constraints and variables. This shows the result holds up to step $t$, finishing the proof. ☐

**Corollary 1.** *Theorem 2's construction extends to an arbitrary matrix $A$ by adding four times more constraints and variables.*

*Proof.* Let $A^+$, $A^-$, $v^+$, and $v^-$ be the absolute values of the entries matching the displayed sign and have zeros everywhere else. We will represent the multiplication $x = Av$ with four variable vectors $x^{++}$, $x^{-+}$, $x^{+-}$, and $x^{--}$ that represent the four ways the signed-components of $A$ and $v$ interact. To do this, we constrain each component with $\Gamma$ using the nonnegative case's construction on the following matrix and vector pairs:

- $x^{++} = A^+ v^+$,
- $x^{+-} = A^+ v^-$,
- $x^{-+} = A^- v^+$,
- $x^{--} = A^- v^-$.

With these constraints on individual components, we have jointly that $x = A_\Gamma^+(v_\Gamma^+ - v_\Gamma^-) + A_\Gamma^-(v_\Gamma^+ - v_\Gamma^-) = A_\Gamma v_\Gamma$. Thus, we can proceed to build the circuit precisely as before. Note this requires four instances of the nonnegative construction, which falls within the claimed bound. ☐
